# Gender-biased perceptions of important ecology articles

**DOI:** 10.1101/219824

**Authors:** Corey J. A. Bradshaw, Franck Courchamp

## Abstract

Gender bias is still unfortunately rife in the sciences, and men co-author most articles (> 70%) in ecology. Whether ecologists subconsciously rate the quality of their peers’ work more favourably if men are the dominant co-authors is still unclear. To test this hypothesis, we examined how expert ecologists ranked important ecology articles based on a previously compiled list. Women proposed articles with a higher average proportion of women co-authors (0.18) than did men proposers (0.07). For the 100 top-ranked articles, women voters placed more emphasis on articles co-authored by women (0.06) than did men (0.02). However, women voters were still biased because they ranked men-dominated articles more highly, albeit not by as much as men did. This effect disappeared after testing read-only articles. This indicates a persistent, subconscious bias that men-dominated articles are considered to be of higher quality before actual assessment. We add that ecologists need to examine their own subconscious biases when appointing students, hiring staff, and choosing colleagues with whom to publish.

## Introduction

Despite a general reduction in gender disparities within academia over time^1–3^, there remains ample gender-bias across scientific disciplines. Experimental evidence shows that scientists tend to rate writings authored by men higher than those authored by women^4^, and that academic scientists tend to favour men applicants over women for student positions^5^. In the United Kingdom, there is also evidence that women academics in science, engineering, and mathematics have more administrative duties on average than men, and hence, less time to do research^6^. Women scientists there also have fewer opportunities for career development and training, and tend to earn lower salaries, hold fewer senior roles, and are less likely to be granted permanent positions than men^6,7^.

Gender bias — in its myriad forms of expression and consequences — is also likely to vary among scientific disciplines. In ecology, despite undergraduates and young researchers having gender ratios closer to parity (as is now the case in most science disciplines^8,9^), senior academic positions in ecology and evolution are still dominated by men^10^. This means that most ecology papers are written by men; for example, in a study examining the proportion of women authorships in papers published from 1990-2011 across 21 science and humanities disciplines, *ecology and evolution* had the seventh lowest proportion of women authors (22.76% of 279012 total authorships)^3^. Women scientists are also consistently under-represented in ecology textbooks compared to baseline assumptions of no bias^9^.

Scientific journals also tend to appoint more men than women on their editorial boards, and editors tend to select reviewers of the same gender as themselves (known as homophily)^11^. Ecologists are also guilty of homophily; for example, men editors selected < 25% women reviewers, but women editors consistently selected between 30 to 35% women reviewers for all papers submitted to the journal *Functional Ecology* from January 2004 to June 2014^12^. Yet, this is not due to the actual performance of women reviewers, because reviewer scores for that journal did not differ between men and women reviewers, and the proportion of papers rejected did not differ between women and men editors^12^. However, there are gender differences in *how* papers are reviewed. For example, from a much broader sample of journals in ecology and evolution, a survey of 1334 ecologists and evolutionary biologists identified that women took longer to review papers than men, and women reviewed fewer manuscripts on average (a logical outcome of being asked less frequently than men to review). In seeming contradiction to the lack of a gender difference in reviewer scores for *Functional Ecology*^12^, men from the broader sample recommended rejection more frequently than did women^13^.

Ecologists can take some heart in the observation that there is little evidence for gender bias in acceptance or citation rates of their papers. In one regional ecology journal (*New Zealand Journal of Ecology*), publication success between 2003 and 2012 was not related to the gender of the authors or that of the editor, but like *Functional Ecology*, editors selected more men reviewers^14^, likely because there are simply more men ecologists from which to choose reviewers. Likewise, there was no author gender bias in citation rate for 5883 ecology articles published between 1997 and 2004 (from the journals *Animal Behaviour*, *Behavioral Ecology*, *Behavioral Ecology and Sociobiology*, *Biological Conservation, Journal of Biogeography, Landscape Ecology*)^15^. A similar conclusion was reached for 507 ecology and evolution articles from five ‘leading’ (but unidentified) journals^10^. Nor was there an overall difference in the acceptance rates of papers according to gender for 2550 ecology and evolution articles (even for single-authored papers), although this differed among journals^10^. However, in one journal (*Behavioral Ecology*), the number of women first-authored papers increased following the implementation of double-blind reviews^16^, suggesting that either women were being given harsher treatment during review, or were less likely to submit when their gender could be identified at the time of submission.

Examining the publication output of 187 individual editorial board members of seven ecology and evolution journals, women had a lower mean *h*-index than did men (after controlling for scientific ‘age’)^17^. Given the lack of evidence for gender bias in citation rates in ecology^10,14,15,18^, it is thought that this was mainly a result of the lower average publication output of women ecologists^17^. Indeed, in a sample of 39 women and 129 men in evolutionary biology and ecology from the same approximate cohort (who held research and faculty positions in the life sciences departments of British and Australian universities), men produced almost 40% more papers than did women, and this difference appeared as early as two years from initial publication^19^. Likewise, a sample of 182 academic biologists (69 women and 113 men) with at least ten years of experience in academia indicated that women produced between 19 and 29% fewer papers after ten years of employment than did men^20^.

Gender differences in publication frequency can occur for many reasons, including possibly having less time to do research^6^, higher demands of motherhood^21–23^, a lower relative tendency compared to men to seek self-promotion^24–26^, fewer academic grants and accolades^27–29^, among other reasons^9,30,31^. Despite no strong evidence yet for gender biases in citation rates in ecology, it is still unclear whether established ecologists — both women and men — subconsciously rate the quality of their peers’ work more favourably than if men are the dominant co-authors, as has been shown for postgraduate students enrolled in communication programs^4^. To test this hypothesis, we have recently compiled a unique dataset to determine which ecology articles are most recommended by ecology experts^32^. From this list of most-recommended articles, we compiled the gender of both the proposers and voters of the articles, as well as the gender of each co-author of the articles themselves (including the gender of the lead author). Specifically, we asked whether ecologists of both genders were swayed by their learned perceptions of article ‘quality’ outside of the review process in terms of: (i) whether men and women proposed or voted for articles more often if they had a higher proportion of men co-authors, and (ii) if there was a correlation between the proportion of women co-authors on an article and its mean rank (as measured and reported previously — see *Material and methods*^32^).

## Results

### Editor pool

The overall proportion of women among the 665 editors we originally contacted to propose articles was 22.1% (i.e., 141 women, 524 men); of these, 14 women (10.0%) and 137 men (26.1%) responded, and 12 women (8.5%) and 101 men (19.3%) proposed articles. These show that men were more likely to respond and propose articles than were women.

### Proposed articles and voting differences

The proportion of women co-authors on the articles proposed by men were on average lower (0.06 to 0.09; mean = 0.07) than those proposed by women (0.13 to 0.27; mean = 0.20), although the data were highly skewed and most proposed articles (77%) had no women co-authors at all (Fig. 1a). When we examined the 100 top-ranked articles voted by women or men only, the bias remained: women voters ranked articles in the top 100 that had more women co-authors (0.029 to 0.093 proportion women) than did those voted by men (0.001 to 0.029) (Fig. 1b).

**Figure 1.**
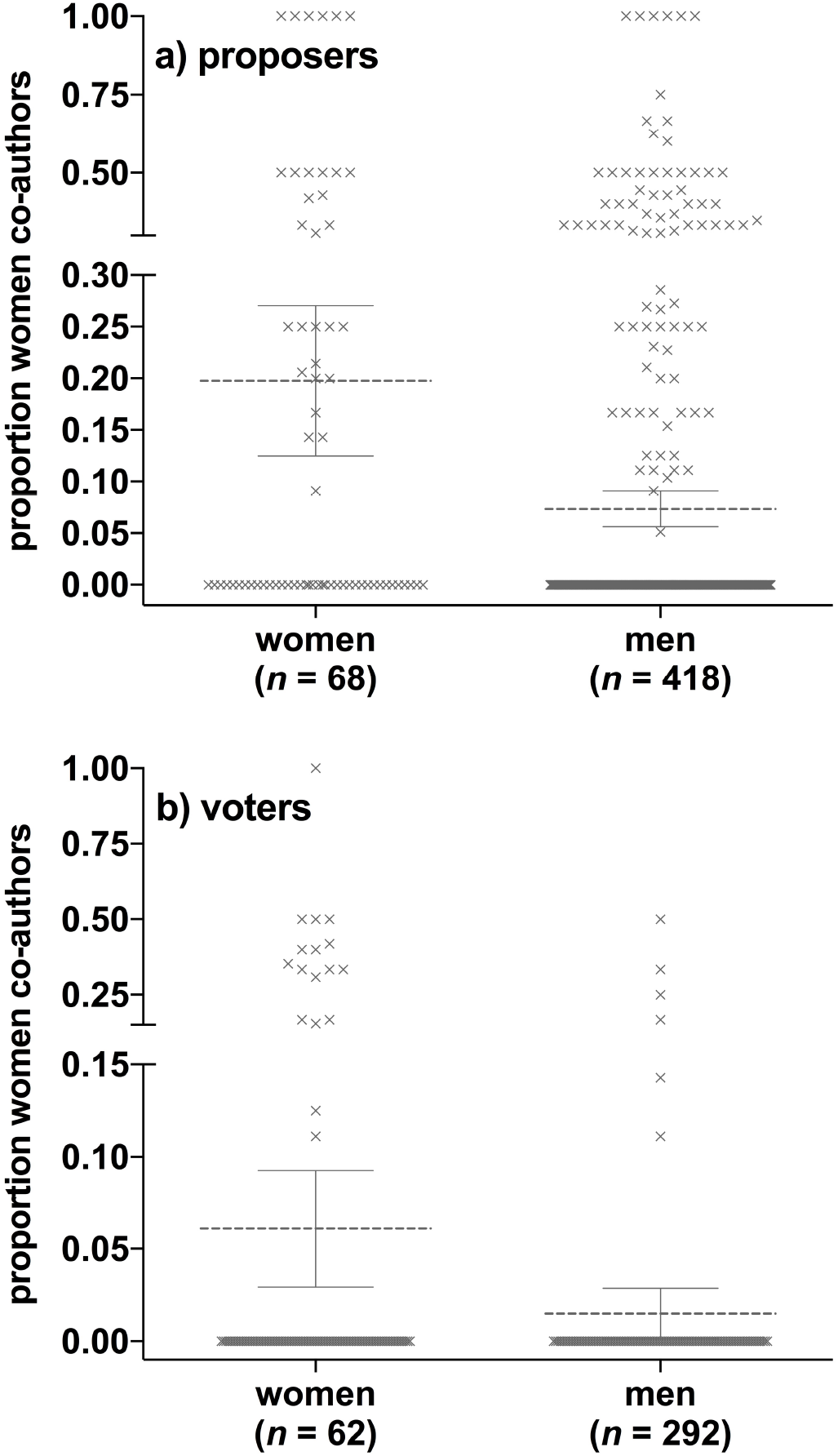
**a**) Mean (dashed horizontal lines) and 95% confidence interval (error bars) of the proportion of women co-authors for the proposed articles relative to the gender of the proposer (articles proposed by 68 women only, and 418 proposed by men only). The values (proportion women co-authors) are ‘scattered’ to show their distribution within each proposer gender; note that 55.9% and 80.1% of the articles proposed by women only and men only, respectively, had no women co-authors (i.e., zero values). **b**) Mean (dashed horizontal lines) and 95% confidence interval (error bars) of the proportion of women co-authors of the 100 top-ranked articles relative to the gender of the voter (62 women and 292 men voted in total). The values (proportion women co-authors) are ‘scattered’ to show their distribution within each voter gender; note that 83% and 94% of the articles proposed by women and men, respectively had no women co-authors (i.e., zero values).

However, even for women voters, there was a tendency to rank men-dominated articles more highly. For women voters only, there was a weak (*β* = 0.03), but non-random (*p*_ran_ = 0.011) correlation between the proportion of women co-authors and the article’s score (from the voting), such that the lower the proportion of women co-authors, the higher they were ranked by women (Fig. 2a). For men voters only, the relationship was stronger (*β* = 0.11) and also non-random (*p*_ran_ < 0.0001) (Fig. 2b). The inverse-score-weighted mean proportion of women co-authors (Σ*w_i_*/*s_i_* = for *i* voters, where *s* = score from 1 to 4, and *w* = proportion of women co-authors) was 0.0277 for women voters, and 0.0251 for men voters (Fig. 2a,b).

**Figure 2.**
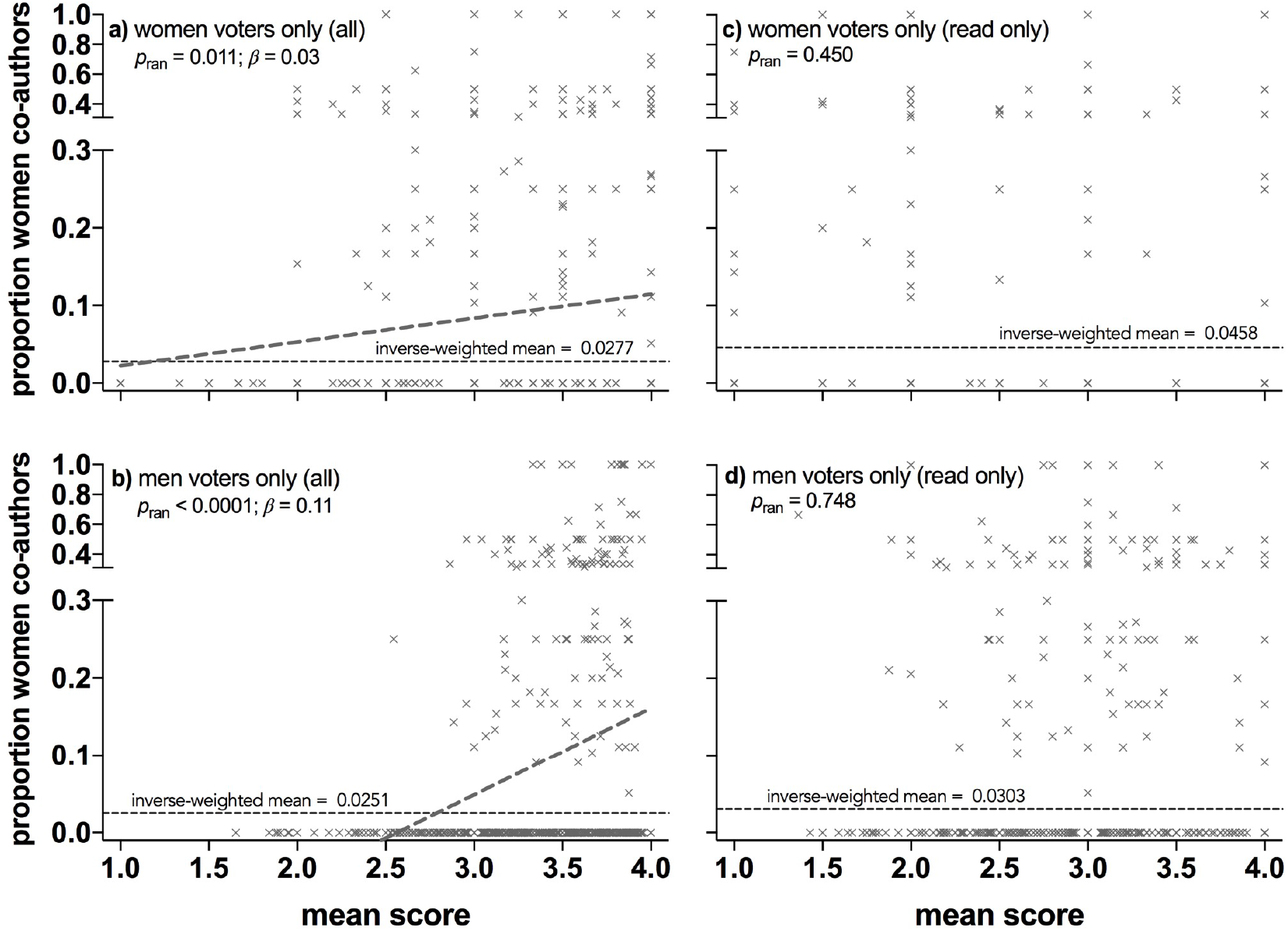
**a**) Proportion of women co-authors on articles relative to their mean rank (score; where lower scores indicate a higher ranking) when voters were restricted to women. There was a weak (*β* = 0.03), but non-random (*p*_ran_ = 0.011) correlation between article gender ratio and score, such that the lower the proportion of women co-authors, the higher they were ranked by women. **b**) Proportion of women co-authors on articles relative to their mean rank when voters were restricted to men. There was a stronger (*β* = 0.11) and non-random (*p*_ran_ < 0.0001) correlation between article gender ratio and score, such that the lower the proportion of women co-authors, the higher they were ranked by men. Also shown in both panels is the inverse-score-weighted mean proportion of women co-authors (Σ*w_i_*/*s_i_* = 0.0277 for *i* women voters, or 0.0251 for *i* men voters, where *s* = score from 1 to 4, and *w* = proportion of women co-authors). **c**) As in a, but when the scored articles were only those actually read by the voters^32^. **d**) As in **b**, but when the scored articles were only those actually read by the voters. The inverse-score-weighted mean proportion of women co-authors for these read-only articles was higher for women-only (0.0458) *versus* men-only voters (0.0303).

### Read-only articles

These non-random relationships could be partially driven by the observation that older articles were more highly ranked than younger articles^32^, and that gender biases in authorship are generally stronger in older articles. So, we also used the ‘read-only’ article scores (in the original survey, voters were asked to indicate whether or not they had in fact read the paper they were scoring; for this new list, article score was unrelated to article age)^32^ for women-only and men-only voters separately. Indeed, the relationships between article rank and proportion of women co-authors disappeared for both women voters (Fig. 2c) and men voters (Fig. 2d), although the inverse-score-weighted mean proportion of women authors was again higher for women voters (0.0458) than men voters (0.0303).

### Lead author

Examining just the gender of the lead author, 510 of the 544 papers (93.8%) proposed had a man as a first author. For the 100 top-ranked papers (read or not), 98 were led by a man; when men alone voted, 99 of the 100 top-ranked papers were led by a man, and when women voted, 96 were. As above, the difference between women and men voters largely disappeared when we examined the read-only list of the 100 top-ranked papers — when women voted, 93 of these was led by a man, and 92 were when men voted.

### Temporal trends

The articles proposed by the entire sample of ecologists indicated a general trend of increasing proportion of women co-authors, from < 5% women co-authors before the 1990s, to the most recent articles published in the last decade exceeding one quarter women co-authorship (Fig. 3).

**Figure 3.**
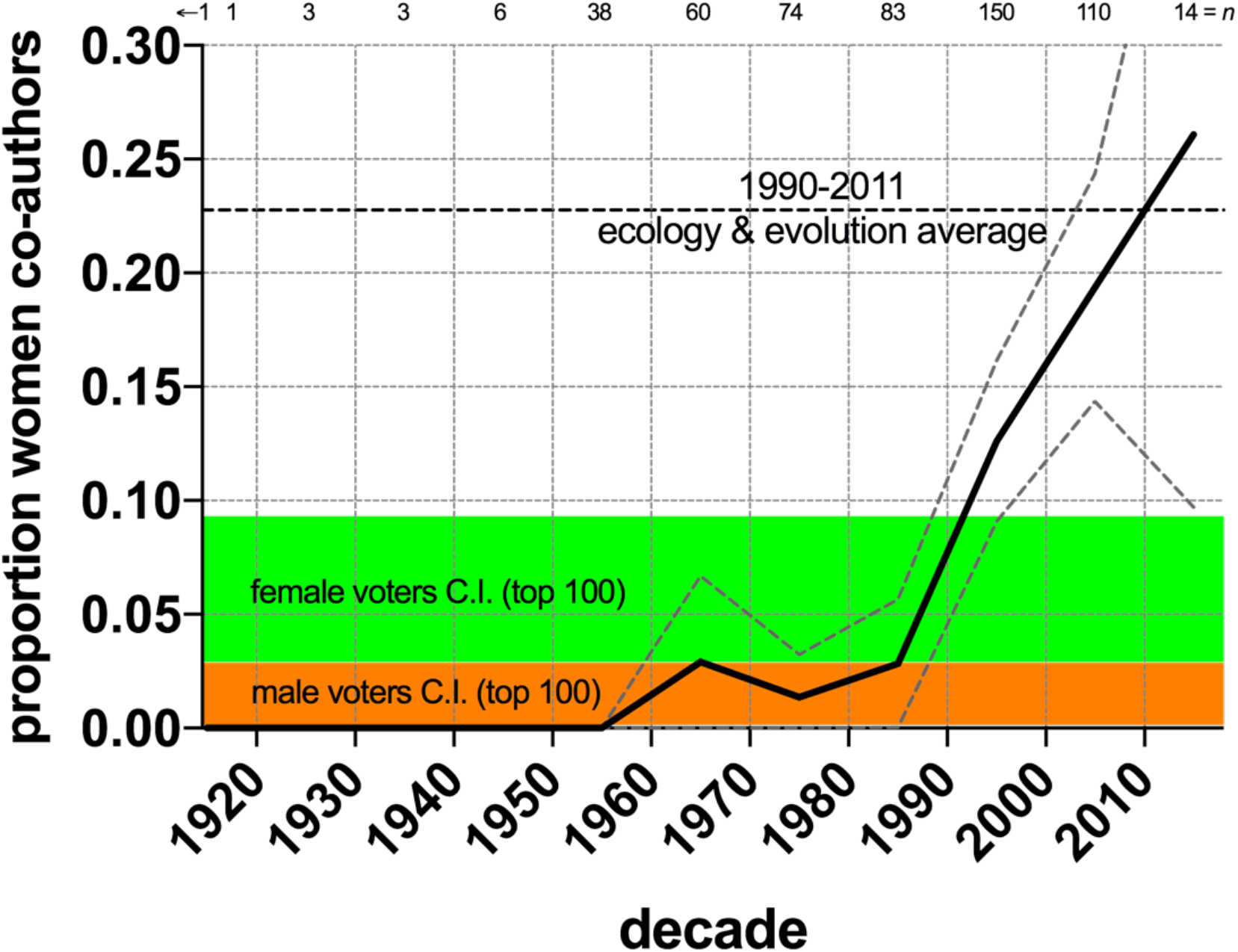
Time series of mean (± 2 standard errors of the mean; grey dashed lines) decadal gender ratio (proportion women co-authors) for all 544 proposed articles. Numbers above the graph indicate sample size (number of articles) used to calculate decadal means (‘←1’ indicates one article from 1858)^37^. For comparison, the proportion of women authorships in articles published from 1990-2011 in ecology and evolution (22.76% of 279012 total authorships; lower black horizontal dashed line)^3^, are shown. Also shown are the 95% confidence limits of the proportion of women co-authors of the 100 top-ranked articles assessed by women (green shaded area: 0.029 to 0.093; Fig. 1b) and men voters (orange-shaded area: 0.001 to 0.029; Fig. 1b).

## Discussion

Our results show that at least for well-established, expert ecologists, both men and women tend to propose and rank articles more highly when they are co-authored by more men, perhaps indicating a degree of homophily when assessing article importance. These results endure despite little evidence that men biologists view themselves as having relatively higher self-perceived expertise than women biologists (according to a sample of 61 men and 190 men tropical biologists)^30^. That article score and the proportion of women co-authors were correlated for both genders can be explained largely by the fact that older papers with which ecologists are at least familiar are generally ranked higher^32^. But because older articles had fewer women co-authors, women ecologists appear to have had little choice but to score the ‘classics’ more highly. Indeed, when we restricted the ranked articles to those that voting ecologists had actually read, the relationship disappeared.

We contend, however, that because assessing the read-only papers demonstrated less of a bias toward men-co-authored papers, this is in fact evidence of a lingering, subconscious gender bias among ecologists. Both men and women ecologists rated articles that they had not actually read higher when they were more men-dominated, yet once they did personally evaluate (read) them, this bias disappeared. This appears to indicate that they had the *a priori* assumption that men-dominated papers would somehow be better. This assumption was stronger in men than women, but it seems that women ecologists are still subject to a persistent form of auto-sexism, perhaps kept flourishing by a remaining academic culture of valuing women’s contributions less than men’s.

This read-only group of younger articles (by 14 years, on average)^32^, combined with the observation that there is an increasing proportion of women co-authors on highly ranked ecology articles, are nonetheless encouraging signs. Indeed, that these highly ranked papers are now (over the last decade) exceeding 25% women co-authors agrees with the approximate overall pool of women co-authors in the general discipline (22.76% women co-authors for articles published from 1990-2011 in ecology and evolution, based on 279012 total authorships)^3^ (Fig. 3).

Despite this increasing trend, our results show that women ecologists are still very much in the minority, both in terms of high-ranking article authorships (i.e., less than one third) and editorships (i.e., less than one fifth). Further, women experts were much less likely to respond to requests to contribute their suggestions of potentially important articles. The underlying reasons for this are unclear, although we hypothesize that it could be explained in part by the observation that expert women ecologists are increasingly and disproportionately requested to take part in surveys, consortia, juries, and committees in an attempt to seek gender parity (e.g., reference^20^). Excessive requests to participate might be exasperating and time-consuming, thus discouraging participation rates relative to men.

Our results highlight two important remaining biases persisting among today’s expert ecologists: (*i*) we all subconsciously bias our opinions of article importance toward those that have at least traditionally been dominated by men co-authors, and (*ii*) men ecologists are still more gender-biased than women ecologists in this regard. While homophily might partially explain these results, it seems apparent that some gender biases against women remain when ecologists assess article quality, and even more so when they judge apparent quality without actually reading the article (i.e., via reputation only). The potential solutions to these problems are varied, including increasing the discussion of the contribution of women ecologists more explicitly in university teaching material^9^, improving flexibility and opportunity in the workplace^2,33^ and at conferences^2,34^ for women, embracing positive discrimination in academic appointments^33^, increasing the prevalence of double-blind reviews^16^, and advocating alternative metrics of citation performance that do not disadvantage women^19^. We further add that all ecologists — especially men — would benefit from serious, personal introspection about their own biases^35^, no matter how uncomfortable an admission of gender bias might be. Denial of one’s own contribution to the problem only serves to perpetuate it^33^. Consciously increasing the number of women ecologists among our students, in our labs, on our editorial boards, requested to review papers, and as co-authors on our manuscripts (something we admittedly failed to do here), will also help to reduce these subconscious biases.

## Methods

The full details of how we generated the list of most recommended ecology articles and how they were ranked are given in Courchamp & Bradshaw^32^; however, we briefly describe the approach and main characteristics of the list here. We contacted the editorial members (*ipso facto*, ecology ‘experts’) of some of the most renowned journals in general ecology: *Ecology Letters, Trends in Ecology and Evolution, Ecology, Oikos, The American Naturalist, Ecology and Evolution* and *Ecography*, as well as all the members of the *Faculty of 1000* Ecology Section (f1000.com/prime/thefaculty/ecol). Of these, we contacted 665 by e-mail to ask them to send us three to five peer-reviewed papers (or more if they wished) that they deemed each postgraduate student in ecology — regardless of their particular topic — should read by the time they finish their dissertation, and that any ecologist should also probably read.

We successfully elicited 147 respondents of the 665 we contacted, who in total nominated 544 different articles to include in the primary list. We then asked these same 665 experts to vote on each of the papers to obtain a ranking, assigning each article to one of four categories: *Top 10, Between 11-25, Between 26-100* or *Not in the top “100”*. We gave one (1) point for each selection of the *Top 10* category, two points for the *Between 11-25*, three points for the *Between 26-100*, and four points for the *Not in the top “100”*. As described in Courchamp & Bradshaw^32^, we averaged all article scores across all randomly sampled sets of votes for each article, and then applied a simple rank to these (ties averaged), thus avoiding any contrived magnitude of the differences between arbitrary score values (i.e., 1 to 4 base scores). The lowest scores therefore indicate the highest ranks.

We manually classified the gender of all proposers, voters, and article co-authors by searching the internet, requesting confirmation from colleagues, or from personal knowledge. We searched meticulously and are confident that we have a correct gender assignment for all people included in the analysis. The Ethics Committee of the *Centre National de Recherche Scientifique* (CNRS, employer of FC) deemed that no ethics approval was necessary for the voluntary and anonymous survey that generated the ranked list of articles.

## Analyses

We took those articles proposed by either women only, or men only, to examine trends between the proposer genders (74% of all proposed papers were proposed only once)^36^. For determining trends between genders of the voters, we subset the entire dataset for women- and men-only voters, tabulating the proportion of women co-authors and the gender of lead authors for the different top-100 ranks resulting from each gender-specific voter subset.

To test for correlations between rankings and gender, we used the proportion of female co-authors for each article as the response variable in all analyses. We also used the same resampling approach in reference^32^ to determine correlations by taking the raw, average scores for each article (independent variable) and compared them to randomised orders of the corresponding correlate (dependent variable) for each test. For each randomised order over 10,000 iterations, we calculated a root mean-squared error (RMSE_random_) and compared this to the observed RMSE between the two variables. When the probability that randomisations produced RMSE ≤ observed RMSE was small (i.e., number of times [RMSE_random_ ≤ RMSE_observed_] ÷ 10,000 iterations ≪ 0.05), we concluded that there was evidence of a correlation.

## Data availability

All code and data for the analysis are available online at github.com/cjabradshaw/HIPE/gender/

## Acknowledgements

We thank the many participating editorial members, as well as to C. Albert and G. M. Luque for help with the survey and article management. FC was supported by BNP-Paribas and ANR grants, and CJAB was supported by BNP-Paris and Australian Research Council grants.

## Author contributions

CJAB and FC conceived and designed the study, and FC collected the data. CJAB did the analyses and wrote the original draft of the manuscript, and CJAB and FC reviewed and edited the manuscript.

